# Length–weight relationships of the French pikes *Esox* spp (Teleostei, Esocidae)

**DOI:** 10.1101/2023.12.01.569518

**Authors:** Lucas Royer, Fabrice Teletchea, Sophie Delavergne, Frédéric Lafitte, Marion Escarpit, Quentin Molina, Emilie Breugnot, Eddy Cosson, Aurélia L’Hostis, Gaël P.J. Denys

**Author notes:** Corresponding author [ ].

## Abstract

Length–weight relationships for Pike *Esox aquitanicus* and *Esox lucius* from France are provided. A total of 9,070 specimens were collected, measured and weighted from 1981 to 2022 throughout France by Departmental Angling Federations and the French Biodiversity Agency during their survey by electrofishing. For all species, the values of *b* are 2.960 for *E. aquitanicus* and 2.987 for *Esox lucius*. We hypothetize this difference by the poor environment where live the Aquitanian pike with no abundant available food and small prey fish species which do not contribute to good conditions for the growth. The study provides the first reference of length–weight relationships for *E. aquitanicus*.

## 1 Introduction

Pikes *Esox* spp. (Actinopterygii, Esociformes) are emblematic fishes because of their strong socioeconomic value for both recreational and commercial fishing (Raat, 1988; Mann, 1996). This genus groups eight species occurring throughout North America and Eurasia (Froese & Pauly, 2023). In France, two species are currently listed: the ubiquitous northern pike *Esox lucius* Linnaeus, 1758 (Fig. 1) and the newly described Aquitanian pike *Esox aquitanicus* Denys, Dettai, Persat, Hautecœur, Keith, 2014 (Fig. 2) (Keith *et al*., 2020). The Aquitanian pike and the allochthonous northern pike co-occur from the Charente to the Adour drainages, because of frequent restocking of the latest since the second part of the twentieth century (Fig. 3) (Denys *et al*., 2014). Both pike species can be distinguished morphologically by their coat coloration patterns, their snout length as well as lateral scales and vertebrae numbers (Denys *et al*., 2014; Jeanroy & Denys, 2019). However, if the northern pike is present in the main rivers and lakes with aquatic vegetation, the endemic species seems to be restricted to small tributaries and coastal catchments qualified as poor environment with sandy substrate and few aquatic vegetation (Fig. 3) (Denys, 2017; Keith *et al*., 2020). *E. aquitanicus* is currently listed as Vulnerable according to the French IUCN Red List of Threatened Species like *E. lucius* because of their risk of hybridization (UICN Comité français *et al*., 2019; Keith *et al*., 2020). As the Aquitaine pike is then a patrimonial and threatened species, riverine managers need tools to apply a good management and conservation policy (Dudgeon *et al*., 2005; Maasri *et al*., 2021). Length–weight relationships (LWRs) constitute primary knowledges used for fish management and stock assessment. They are necessary for estimating fish biomass from sampled length data, as well as fish growth, and are useful for ecological modelling (Froese, 2006). For that, length *L* and weight *W* are related with a mathematical formula *W = aL*^*b*^ including two parameters: a coefficient *a* and the allometric growth parameter *b* (Keys, 1928). These two parameters are essential to understand the growth of each species. Each pike species has at least one LWR published study (e.g., Kapuscinski *et al*., 2007; Verreycken *et al*., 2021; Giannetto *et al*., 2016; Huo *et al*., 2017; Parker *et al*., 2018), except the Aquitanian pike. Irz *et al*. (2022) published also a LWR of “*Esox lucius*” from the ASPE database collecting lengths and weights data of French fish species collected by the French Agency of Biodiversity (OFB) since 1980s, but without distinguishing the two species. Well, working on data from badly identified taxa can induce some bias on their management (Bortolus, 2008).

**Figure 1.**
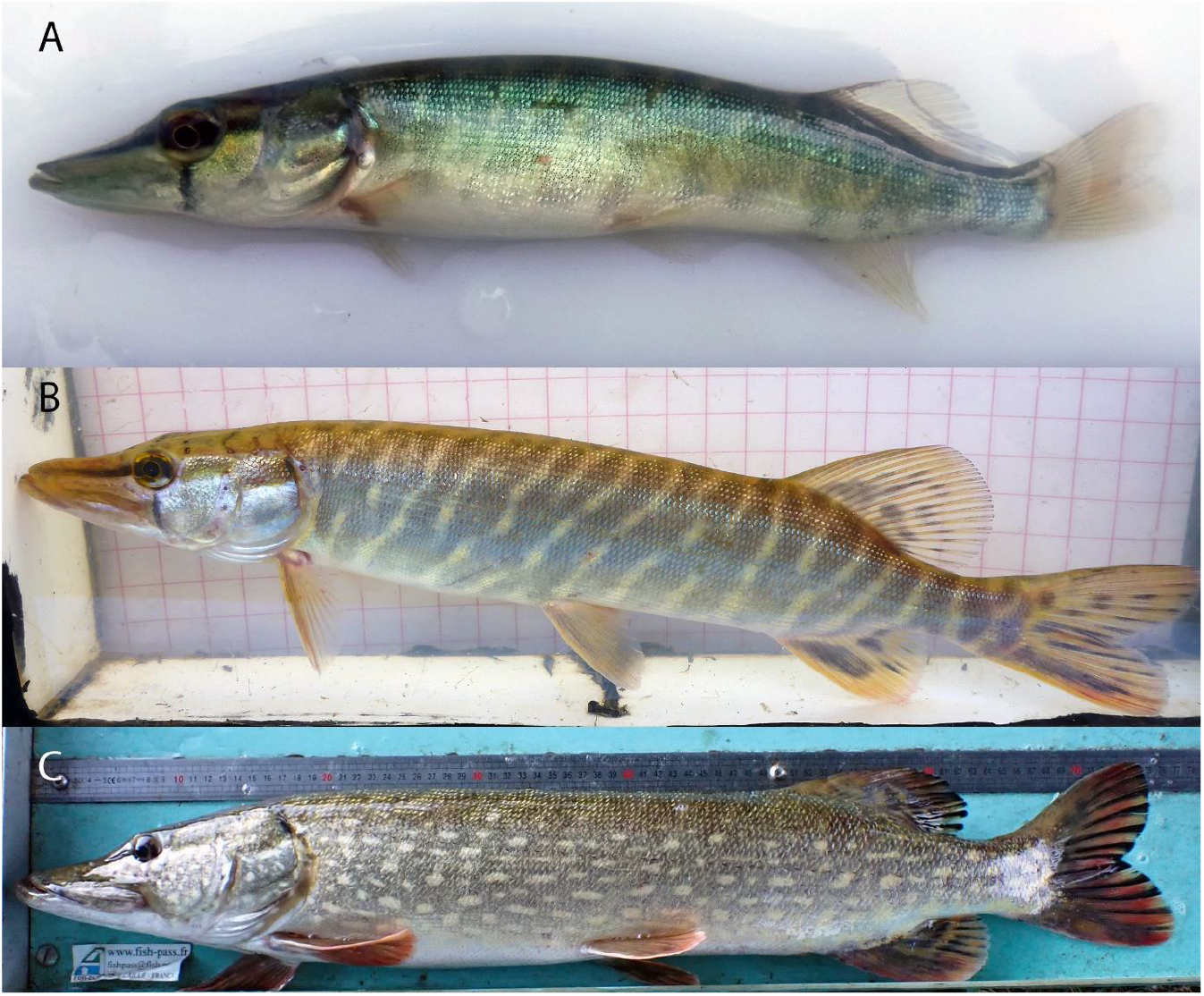
The northern pike *Esox lucius*: 92 mm from the Vie river at Poiré-sur-Vie (A), 278 mm from the Yser river (B), 770 mm from the Oise river (C); credit photos: Hydrosphere, Fishpass, OFB.

**Figure 2.**
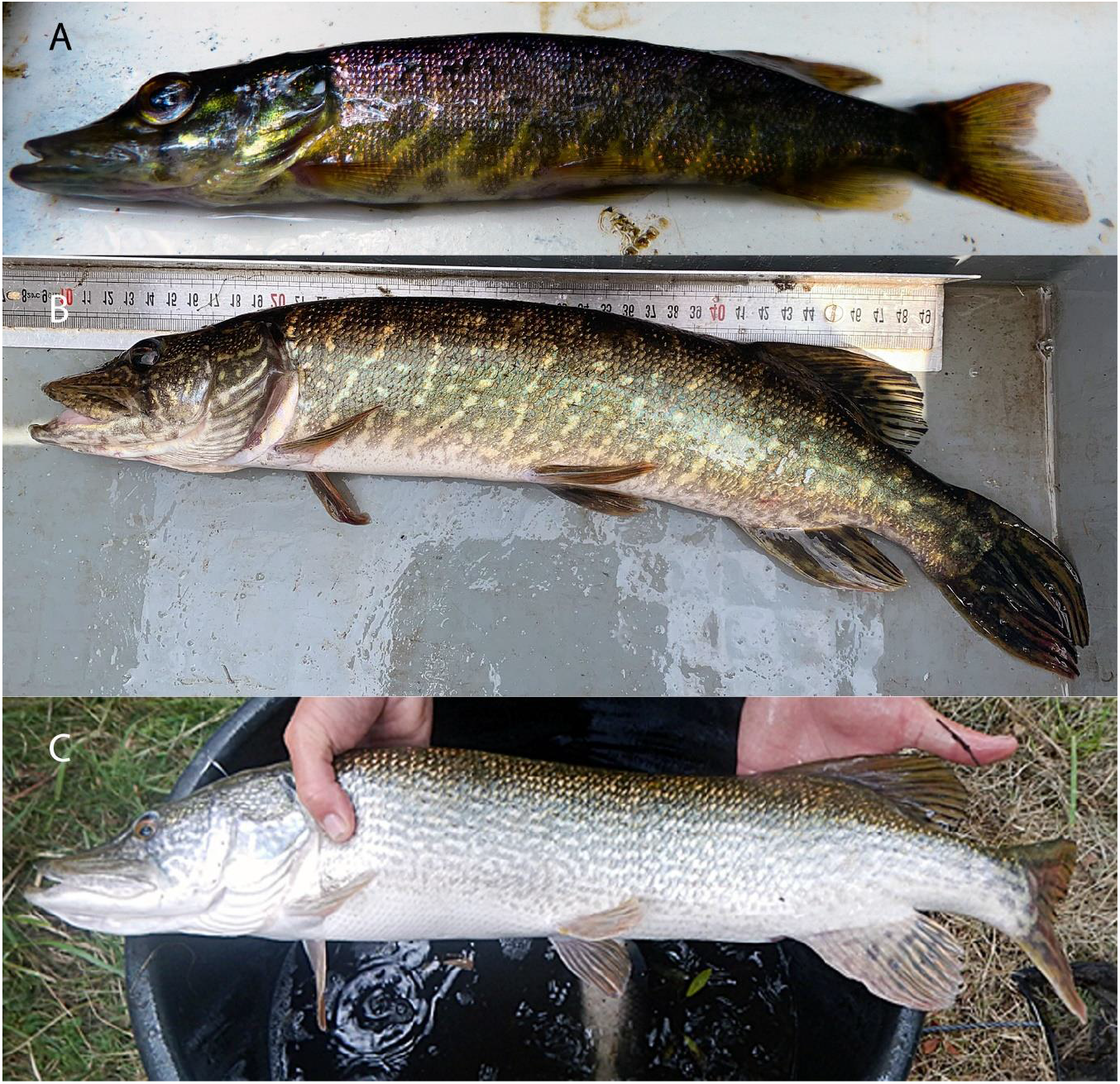
The Aquitanian pike *Esox aquitanicus*: 111 mm for 9 g from the Ciron stream (A), 480 mm for 816g from the Courant mort brook (B); 747 mm for 3,200 g from the Jalle du Sud stream (C); credit photos: FDAAPPMA47, FDAAPPMA40, FDAAPPMA33.

**Figure 3.**
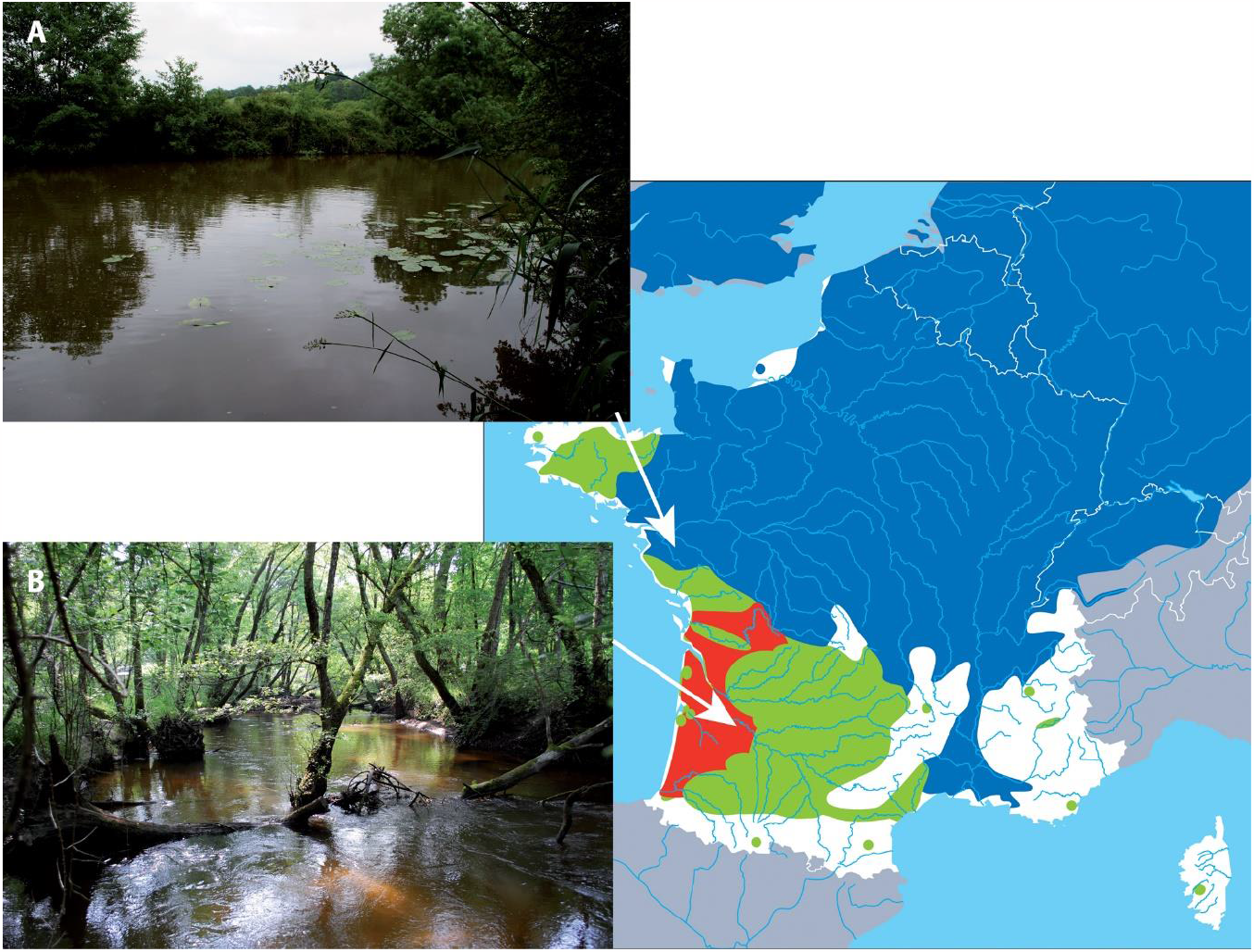
Distribution area of the two pike species in France (*Esox aquitanicus* in red, *Esox lucius* in its native area in blue and introduced in green) (modified and adapted from Keith *et al*., 2020 and Denys, 2017), with examples of typical habitats of both species: the Sèvre nantaise (A) as a large river with many aquatic vegetation and the Ciron stream (B) as watershed with a poor environment (sandy substrate and no aquatic vegetation except stumps); credit photos: G. Denys.

The aim of this study was to provide the first LWR for *E. aquitanicus* based on an extensive sampling throughout its distribution area, to compare it with data of French *E. lucius* in order to know if there are some differences between both species, and to know if misidentifications in the ASPE database of Irz *et al*. (2022) have repercussions on their results.

## 2 Material and methods

Data for Aquitanian pike were collected from 1996 to 2022 during the monitoring of the Departmental Angling Federations and the French Biodiversity Agency (OFB) which aim to make the diagnosis of the physical, trophic and physico-chemical state of the environments in order to best adjust the fish management measures. Specimens were caught by electrofishing inventories campaigns before to be released. Populations previously characterized as hybrids or introgressed by Denys *et al*. (2014, 2018) and Denys (2017) were removed from our dataset as well as the locations already known to have been restocked. Thus, our dataset is composed by only populations already characterized as pure Aquitanian pike according to morphological (Denys *et al*., 2014; Jeanroy & Denys, 2018) or molecular data (Denys *et al*., 2014, 2018; Denys, 2017). Total length (*L* in cm) was measured to the nearest millimeter and weight (*W* in g) was determined with a digital balance to an accuracy of 0.1 g. Additional data (n = 338) from locations already knows to shelter Aquitanian pike without any sympatry with the allochtonous species were added from the ASPE database (Irz *et al*., 2022). The list of locations and measurements are available in Supplementary data 1 and 2.

For northern pike, data were extracted from the ASPE database excluding those from the Adour-Garonne drainages (location codes beginning by “05”) in order to exclude any data corresponding to the Aquitanian pike, and during the period between 1981 to 2017. Only individual measurements were considered. A map synthetizing the sampling is given in Figure 4.

**Figure 4.**
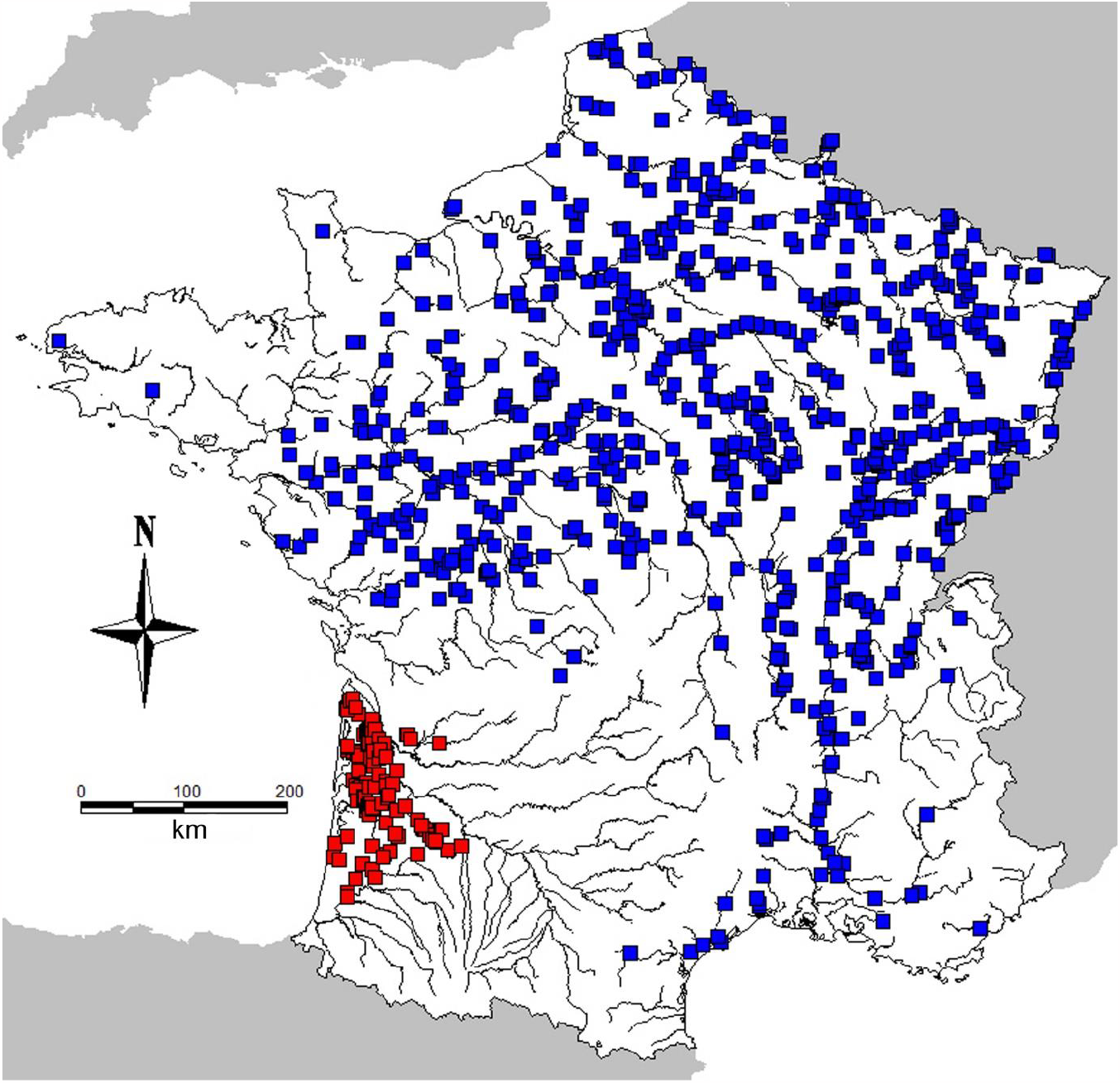
Locations of both pike species (*Esox aquitanicus* in red and *Esox lucius* in blue) from where lengths and weights data came.

In both datasets, records with weights lower or equal to 1 g were removed considering weighing scales used were not sufficiently accurate to have reliable data for these young offsprings. Lengths and weights averages were compared using a *u* test of Mann-Whitney after having checked a non-normal distribution with a Shapiro-Wilk test (p-value < 2.2 e^-16^). Length–weight relationships were calculated following the method of Kuriakose (2014) using the equation log10*W* = log10*a* + *b* log10*L*, where *a* is the intercept on the *Y*-axis of the regression curve and *b* is the regression coefficient (Ricker, 1975). LWRs were calculated for unsexed individuals, as the sex determination is not done by the angling federations nor the French Agency of Biodiversity, because it is not included in their protocol of surveys. The 95% confidence limits of *a* and *b* (CL 95%) were also computed (Froese, 2006) for both equations. Linear regressions and all statistic tests were performed using the R package (R core team, 2022) using lmtest, stats and dplyr packages following Irz (2022). In order to validate the regression model for each dataset, a Rainbow-test was done in order to verify if the residuals mean was equal to 0 (Utts, 1982). Their p-value = 1 and 0.147 for respectively *E. aquitanicus* and *E. lucius* indicated a leverage effect. So, the Cook’s D distances were then calculated and data exceeding 4/N (N being the number of observations) were removed from the dataset (Bollen & Jackman, 1990).

## 3 Results

Data on lengths and weights are provided for a total of 3,657 Aquitanian pikes and 5,413 northern pikes. The average sizes and weights are respectively 10.7 cm for 20.2 g and 30.2 cm for 339.8 g. Both u-tests of Mann-Whitney on lengths and weights indicates that the average data for *E. aquitanicus* is lower than those of *E. lucius* (W_lengths_ = 1341292; W_weights_ = 1413738; p-value < 2.2 e^-16^ for both datasets). The number of specimens, TL range, parameters a and b with their 95% CL and the correlation coefficient (*R*^2^) are reported in Table 1.

**Table 1.** Descriptive statistics and parameters of total length (TL, in cm) – weight (W, in g) regression for the Aquitanian pike (South West of France) and northern pike from France; SD means standard deviation.

| Species | Sex | n | TL range<br>(SD) | W range<br>(SD) | a | Range a (95% CL) | b | Range b (95% CL) | R<sup>2</sup> |
| --- | --- | --- | --- | --- | --- | --- | --- | --- | --- |
| *Esox aquitanicus* | All | 3,657 | 5.0 – 72.0<br>(5.834) | 1.2 – 2,500<br>(85.060) | 0.008 | 0.007–0.008 | 2.960 | 2.947–2.973 | 0.982 |
| *Esox lucius* | All | 5,413 | 5.9 – 125.0<br>(15.355) | 2.0 – 12,450<br>(708.559) | 0.007 | 0.006–0.007 | 2.987 | 2.978–2.995 | 0.988 |

The *R*^2^ values are respectively equal to 0.982 and 0.988 demonstrating a strong correlation between lengths and weights for both species (Fig. 5).

**Figure 5.**
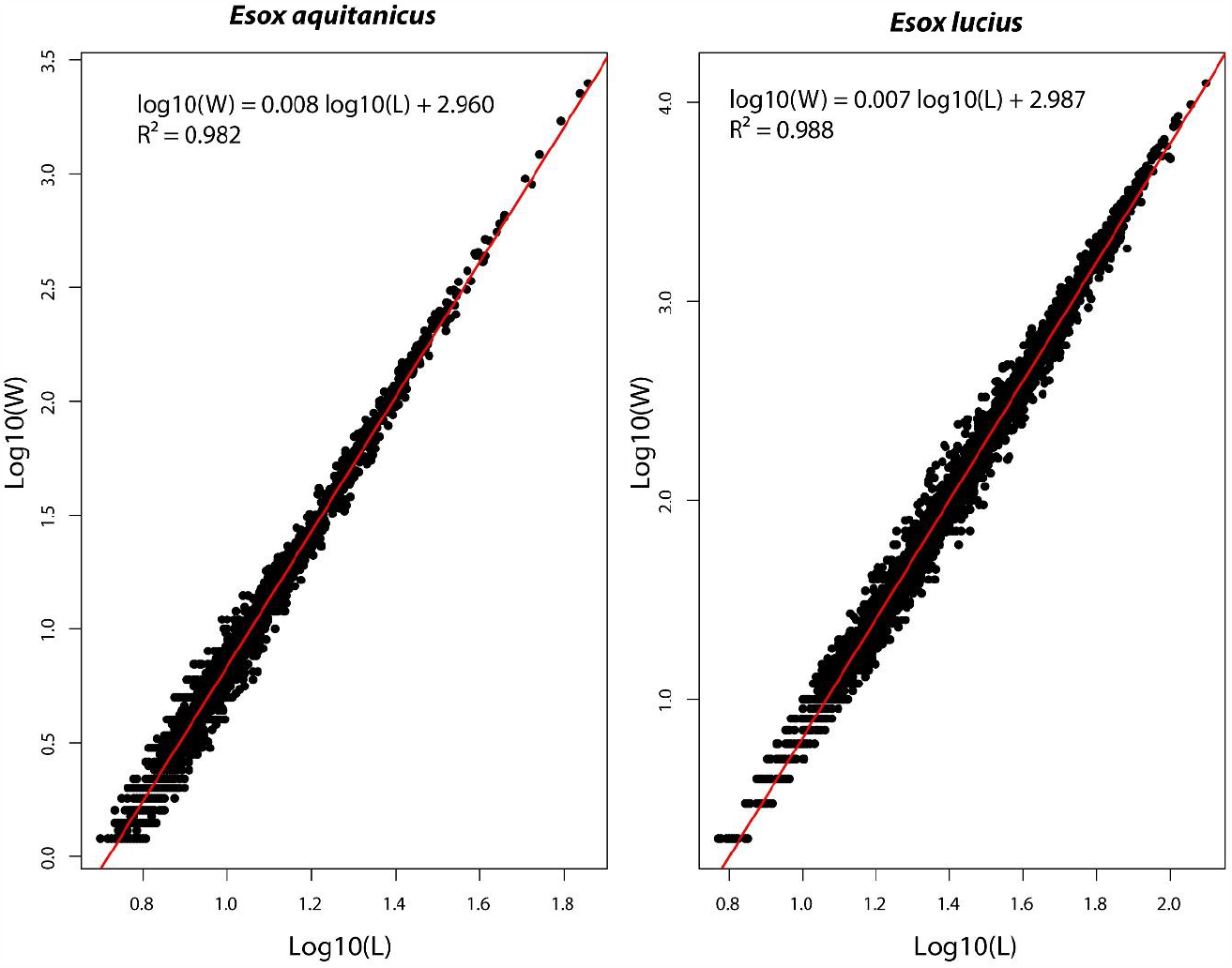
Lengths (L) - weights (W) regressions of *Esox aquitanicus* and *Esox lucius*. Data were log10 transformed.

*E. aquitanicus* has a lower *b* parameter than *E. lucius* (2.960 *vs*. 2.987) as their range do not overlap (Table 1).

## 4 Discussion

LWR of *E. aquitanicus* and French *E. lucius* were established from the analysis of 9,070 fish. Aquitanian pike are smaller than northern pike with a maximum size conserved in our dataset of 72 cm for 2.5 kg. Fish sizes are positively correlated with the surface area of habitat and negatively with the temperature (Denys *et al*., 2016; Kennedy & Rennie, 2023). But Aquitanian pike lives in small tributaries and little coastal basins (Keith *et al*., 2020), and in southwestern France which is one of the warmer regions in the country (e.g., Parey *et al*., 2007). However, anglers accounts and data from bibliographical archives described larger specimens: 85 cm for 5 kg (Lagardère, 2020), 107 cm for 9 kg (Glize, 1993) and large fish from 12 to 15 kg (Cahuzac, 2001). So, when the environmental conditions are favourable, Aquitanian pike can reach comparable sizes than northern pike.

The *a* parameter for the Aquitanian pike is significantly smaller than the one of *E. lucius* (Table 1). Whereas the *b* parameters are respectively 2.960 and 2.987 in accordance with Carlander (1969) assuming that *b* normally falls between 2.5 and 3.5. Values of *b* lower than 3, meaning a “negative” growth, so both pike species become slimmer with increasing length. Comparing both *b* values, the one *E. aquitanicus* is lower than *E. lucius* (Table 1). The hypothesis that *E. aquitanicus* would be more elongated than *E. lucius* would be false because of its shorter snout and the fewer number of vertebrae (Denys *et al*., 2014). So, the northern pike would be heavier than the Aquitanian pike for the same given size. Differences between a and b values may be explained by the poor environments where it lives (sandy substrates, few aquatic vegetation, low biomass) conferring a lower primary productivity (Tales *et al*., 2004). The specific richness is also low (about 5 to 8 co-occurring fish species) (see CGA hydroregion from Santoul *et al*. (2004) and assemblage type 1 from Park *et al*. (2006)), and closed to the historical native ichthyofauna community in the Adour-Garonne basin (Keith *et al*., 2020) with little sized species such as minnows *Phoxinus* spp, stone loaches *Barbatula* spp and gudgeons *Gobio* spp which are not in elevated densities considering the particular habitat (Tales *et al*., 2004). However within pikes, the growth rate is correlated with the size of the eaten preys (Hart & Connellan, 1984) and the quantity of available food (Kozłowski *et al*., 2012; Kennedy & Rennie, 2023). A study on ecological traits of the Aquitanian pike is sorely needed to support this hypothesis.

Our *a* and *b* parameters for *Esox lucius* are exactly the same as given by Irz *et al*. (2022), as well as the *R*^2^ (0.987). Their huge dataset (n = 9,535) in the ASPE database and removing data after the Cook’s D distances calculations has certainly drowned the signal of *E. aquitanicus* in the dataset. The addition of data from angling federations allowed as well as working on populations correctly identified has allowed to bring new knowledges on the endemic species. Angling federations have useful data and naturalist observations which deserve to be known and used in monitoring, ecological studies and conservations programs (Maasri *et al*., 2021).

Their datasets are then complementary with the database ASPE. However, the data centralization from each departmental angling federation and the access is still a huge challenge. Managers are encouraged to integrate in their protocol the sex determination. This kind of data would allow to highlight a potential sexual size dimorphism already known within pike species (e.g., Craig, 1996) with female growing faster than males. Nevertheless, these cases are correlated with the availability of food resources and cool temperatures (Kennedy & Rennie, 2023). But the geographical context does not fulfil these conditions.

Finally, our study provides then the first data of LWR for Aquitanian pike which could be implemented in FishBase (Froese & Pauly, 2023). This LWR could be useful for managers for estimating weights from measurements and to know if the environment of a location brings ideal conditions for growth, as well as for other disciplines in biology.

## Acknowledgements

This work was supported by the Muséum national d’Histoire naturelle (MNHN), the Unit PatriNat 2006, the UMR BOREA 8067, the Lorraine University, the French Biodiversity Agency and the Union of Angling Federations of the Adour-Garonne Basin (UFBAG). This study is included within a global project about the Aquitanian pike BIOESOX funded by the UFBAG and the French Federation of Angling in France (FNPF). We thank all people from the Angling departmental federations and the OFB for providing data.

## Figure legends

Supplementary data 1. List of locations of *Esox aquitanicus* used in this study.

Supplementary data 2. Measurements of *Esox aquitanicus* used in this study.

## Notes

### Competing Interest Statement

The authors have declared no competing interest.

